# ZNF423 orthologs are highly constrained in vertebrates but show domain-level plasticity across invertebrate lineages

**DOI:** 10.1101/2020.03.09.984518

**Authors:** Bruce A. Hamilton

## Abstract

*ZNF423* encodes 30 C2H2 zinc fingers that bind DNA and a variety of lineage- and signal-dependent transcription factors. *ZNF423* genetic variants are proposed to cause neurodevelopmental and ciliopathy-related disorders in humans. Mouse models show midline brain defects, including cerebellar vermis hypoplasia, and defects in adipogenesis. Here I show strong protein sequence constraint among 165 vertebrate orthologs. In contrast, orthologs from invertebrate lineages, spanning larger time intervals, show substantial differences in zinc finger number, arrangement, and identity. A terminal zinc finger cluster common among other lineages was independently lost in vertebrates and insects. Surprisingly, a moderately-constrained non-C2H2 sequence with potential to form a C4-class zinc finger is a previously-unrecognized conserved feature of nearly all identified homologs. These results highlight evolutionary dynamics of a likely signal integration node across species with distinct developmental strategies and body plans. Functions of the newly identified C4-like sequence and lineage-specific fingers remain to be studied.

## Introduction

*ZNF423* and its paralog, *ZNF521*, each encode 30 C2H2 fingers, fourth-most among curated human proteins [1]. These paralogs arose early in the vertebrate radiation (apparent single copy in hagfish) and are readily distinguished among vertebrate genomes by characteristic sequence differences. C2H2 zinc fingers (ZFs) are a widely-distributed sequence motif that imparts structural specificity useful in binding defined targets [2-4], often through clusters of three or more fingers per target. C2H2 ZF structure is mediated by the four zinc coordinating residues for which it is named and three hydrophobic residues that form the core, all with defined spacing. Other ZF classes, especially C4 have less sequence homology and are therefore harder to identify from sequence alone [5]. While perhaps best known as a sequence-specific DNA-binding domain, zinc fingers can also bind specific RNA [6, 7] and protein [8, 9] targets, showing versatility and adaptability of this structural fold family [10-13].

Adjacent clusters of C2H2 ZFs in ZNF423 act as binding modules, first shown by Tsai and Reed in the 1990s. ZF28-30 were first identified as binding to lineage-restricted Early B-cell Factor (EBF) family factors in rat olfactory neurogenesis, retarding lineage differentiation by inhibiting the formation of EBF:EBF dimers [14]. ZF2-8 showed high-affinity DNA binding to inverted GCACCC repeats as an apparent dimer or mulitmer, with weak binding to a single half-site [15]. In the same study, multimerization was shown to require either ZF28-30 or a non-motif sequence between ZF25 and ZF26. Hata and colleagues showed that internal clusters mediated binding to Bone Morphogenetic Protein (BMP) signaling-dependent SMAD proteins (ZF14-19) and the BMP response element (ZF9-13) in frog embryos and cultured mammalian cells [16]. During DNA damage response ZF3-8, overlapping the DNA binding domain, bound centrosomal protein CEP290, while ZF11-23, covering the BMP response element and SMAD-interacting fingers, bound PARP in human cells [17, 18]. ZNF423 bound with retinoic acid receptors [19] in neuroblastoma cells (and ZNF423 appeared to be a critical factor for retinoic acid signaling in cortical brain development [20]) and with Notch intracellular domain [21], although these binding sites have not been mapped. A consistent feature of these studies has been that ZNF423 binding partners downstream from signaling pathways (SMADs, RAR, and Notch intracellular domain) antagonized ZNF423:EBF heterodimers, suggesting a mechanism to make EBF-mediated lineage programs responsive to extracellular cues.

Genetic evidence has demonstrated a substantial range of *ZNF423* phenotypes. Loss of function variants were strongly depleted from human non-disease populations [22, 23] and putative rare mutations have been proposed for patients with neurodevelopmental disorders [17, 24]. In mice, null and reduced-expression alleles caused midline hypoplasia in cerebellum and forebrain [25-27] and showed defects in adipogenesis [28, 29] and wound healing [30]. In mouse cerebellum, ZNF423 was required for proliferative response to SHH [31]. The extent of brain malformation was influenced by strain background, consistent with the idea that ZNF423 coordinates cellular responses across multiple developmental signals [32, 33] and included hindbrain choroid plexus [25, 34]. Mice engineered to lack EBF-binding and SMAD-binding domains had hypoplastic defects less severe than null and with different patterns of neurogenic defects, albeit with small sample size [35]. New work from our group showed mild midline abnormalities in mice lacking ZF1 or ZF25 and an adjacent putative C4-like zinc finger, neither of which yet has a known interaction partner. We saw more substantial partial loss of function phenotypes for mice lacking ZF15-18 [36]. Surprisingly, we found no evidence for structural brain abnormality in mice lacking ZF12, in the annotated BMP response element-binding domain. Consistent with its developmental role in cellular differentiation, ZNF423 has also been implicated as a modulatory factor in neuroblastoma [19] and glioma [37], where patient samples with higher ZNF423 expression correlated with better survival. A Drosophila homolog, DmOAZ, is expressed in nervous system and filzkörper, with structural abnormalities noted in the latter tissue in presumptive null mutations [38].

Results described here address a gap in understanding evolutionary constraints among ZNF423 homologs across different time scales. The results showed strong constraint in amino acid sequence among 165 diverse vertebrate genomes, but plasticity in both sequence and domain arrangements across larger taxonomical divisions. Specifically, invertebrate lineages showed changes in ZF numbers both within identifiable clusters and added carboxyterminal (C-terminal) to ZF domains homologous to vertebrates. This analysis also identified a novel CxxC-x(10-31)-CxxC motif, reminiscent of treble clef fold group zinc fingers but not annotated by common motif algorithms. This C4-like feature between ZF25 and ZF26 of vertebrates was a conserved feature of ZNF423 homologs across bilateria.

## Results

### *ZNF423* orthologs are highly constrained among vertebrates

To examine levels of constraint among vertebrate ZNF423 orthologs, homologs that distinguished ZNF423 from ZNF521 were manually curated from multiple sources, including general and taxonomy-limited BLAST searches and publicly curated orthologs. Excluding duplicate species from a single genus, sequences labeled low-quality in their accession record, and sequences that appeared fragmentary, this produced 165 distinct sequences, spanning a wide range of jawed vertebrates and one hagfish (Supplemental Table S1) covering ∼615 million years since the last common ancestor [39].

The vertebrate alignment ignored the first two short coding exons because annotated starting peptides were variable across species, likely due to incomplete genome assemblies or gene models. The starting peptide MSRRKQ found in mouse (NP_201584.2) and rat (NP_446035.2) was widely distributed in all vertebrate lineages, and among invertebrate homologs. The starting peptide MHKK in the human reference sequence (NP_055884.2) was not found outside of catarrhine primates (apes and Old World monkeys) and was internal to an MSRRKQ start where both peptides were found. The first two coding exons (33 codons in mouse, 25 in human) were relatively short, did not encode a zinc finger, and were missing from 67 (and questionable in 8 more) of the 165 inferred orthologs. The analysis pipeline therefore trimmed each sequence to the aligned position of a moderately conserved methionine in exon 3 that was aminoterminal (N-terminal) to the first C2H2 zinc finger and the most N-terminal residue present across all 165 sequences (M61 in human, V69 in mouse).

Vertebrate alignments showed a remarkable degree of constraint throughout the remaining protein sequence. Choice of alignment algorithms and parameters made little difference. Default parameters in MUSCLE were used for the analysis shown. Alignment gaps occurred only between zinc fingers, typically between clusters of functionally related zinc fingers defined by previous experimental data. Of 1224 sites aligned from the human reference sequence, 609 (49.8%) were invariant. To visualize constraints, substitutions were scored in MAPP [40], which considers both phylogenetic structure of the samples and physicochemical properties of substituted residues (Figure 1A). Similar graphs were obtained using alternative approaches, including dN/dS ratio, ScoreCons [41] and ConSurf [42, 43] (Supplementary Figure S1). This analysis predicted additional functional constraint among vertebrates on non-motif sequences between ZFs 1 and 2, 8 and 9, 13 and 14, and 25 and 26.

**Figure 1.**
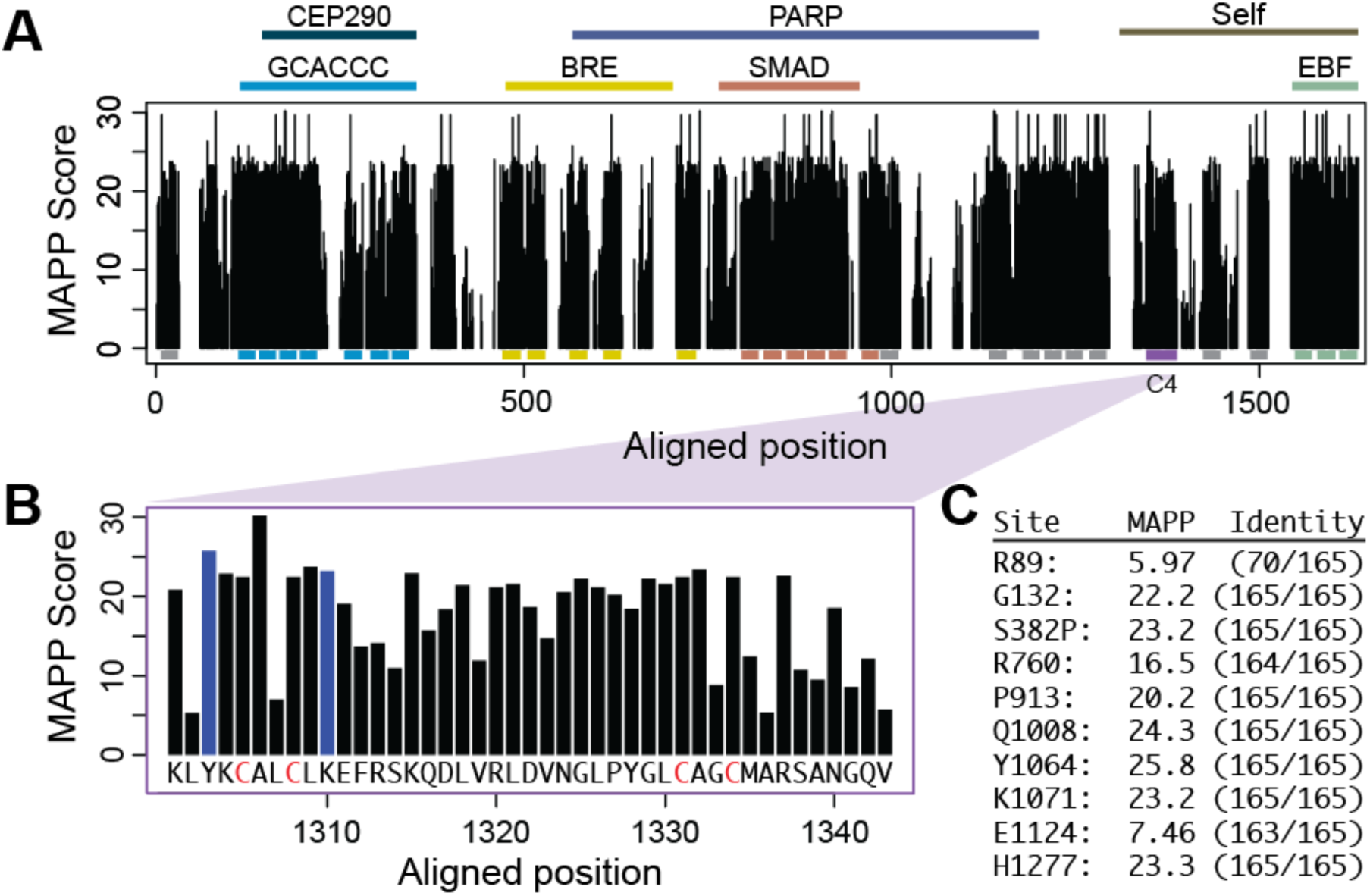
ZNF423 domain architecture and vertebrate sequence constraint. (A) MAPP scores by alignment position for 165 vertebrate sequences in Supplemental Table S1. Higher values indicate stronger constraint, but note that different residues have different maximal scores depending on physicochemical properties. Reported binding regions are indicated above the graph for EBF and SMAD transcription factors, the BMP response element (BRE), the GCACC consensus DNA sequence, centrosomal protein CEP290, and Poly (ADP-ribose) polymerase (PARP). Locations encoding individual zinc fingers are indicated below the graph, colored by interaction. (B) MAPP scores and human amino acid sequence for a conserved potential C4 zinc finger. All four cysteine residues are present in all species. Y1064 and K1071 in the human reference sequence are indicated in blue. (C) MAPP score and fraction identity for 10 human variant sites studied by Deshpande et al. [36].

Of the three constrained non-motif sequences, only the sequence between C2H2 ZF25 and ZF26 was conserved with invertebrate homologs. Among vertebrates, this region contained an invariant YxCAxCLK-(x14)-GxPxGxCxxC sequence, which has potential to form a C4-like zinc finger or treble clef-like fold (Figure 1B). Interestingly, this overlaps sequence implicated in multimerization of ZNF423 [15] and includes two of ten clinical variants of uncertain significance tested in mice [36]. Analysis of the 165 vertebrate orthologs showed that these two sites (Y1064 and K1071 relative to human RefSeq NP_055884.2) and seven of the ten sites overall were invariant across all 165 available vertebrate sequences (Figure 1C).

**Supplementary Figure S1.**
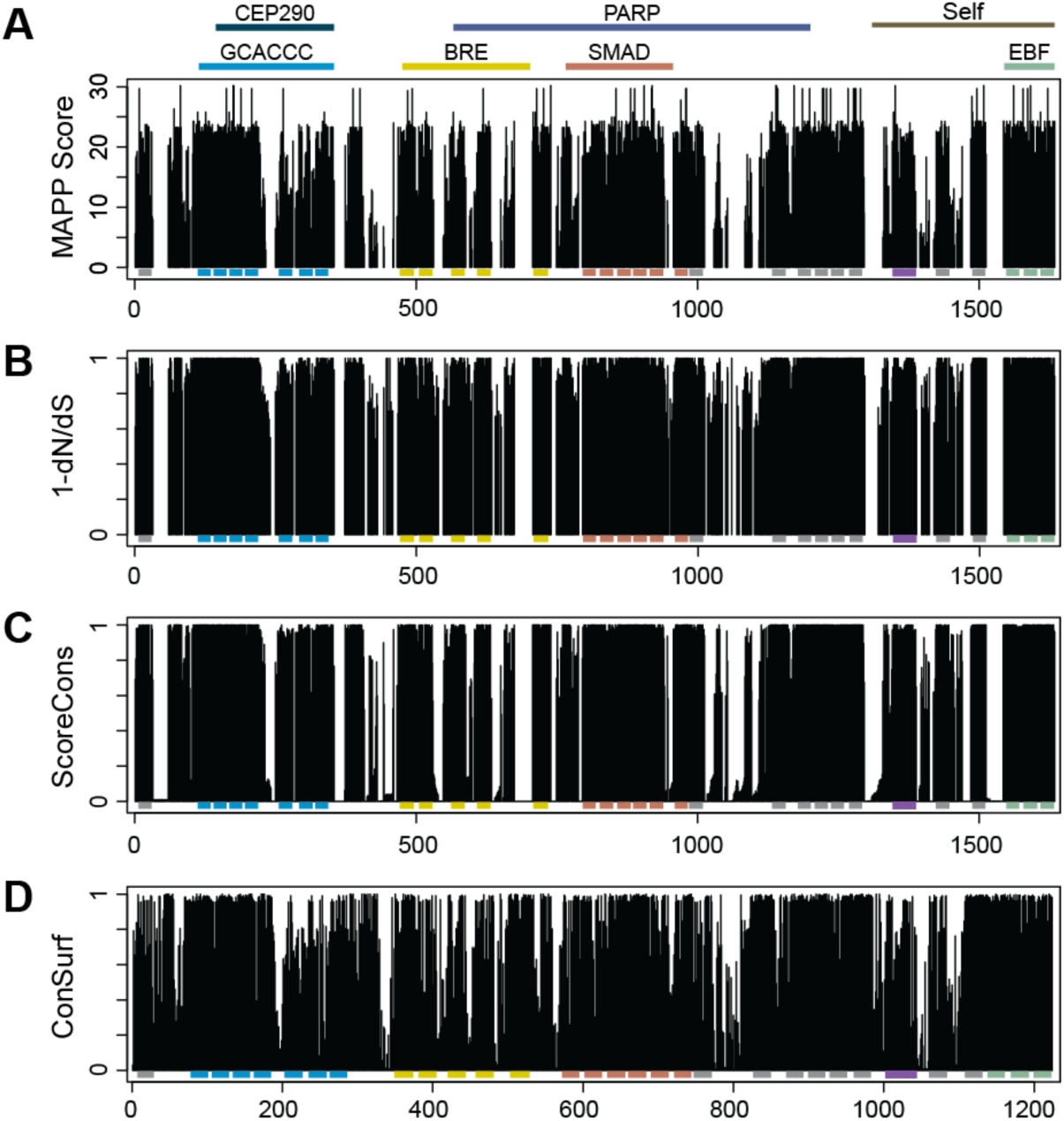
General features of vertebrate ZNF423 constraint are robust to method. (A) Median substitution score from MAPP repeated from Figure 1. (B) 1-dN/dS for each codon shows strong evidence for negative selection throughout the coding sequence. dN/dS was <1 for all sites. (C) ScoreCons normalized measure on MAFFT-alignment of the same sequences. (C) ConSurf analysis, plotting normalized e^-score^ to produce positive values from 0 to 1, indexed to human ZNF423 as the reference sample. Zinc finger motifs are indicated by bars below each plot.

### Invertebrate homologs show ZF gain and loss across longer time scales

Public annotations and iterative BLAST searches of public databases identified 76 unique invertebrate homologs, after removing sequences annotated as partial or low-quality and sequences substantially shorter than others in the same phylogenetic group (Supplemental Table S2). Homologs were single-copy per species and presumed orthologs to both ZNF423 and ZNF521. Homologs were present in all major bilaterian lineages, including Arthropods, Brachiopods, Echinoderms, Hemichordates, Mollusks, and Nematodes. Each of these lineages included examples with the MSSRKQ N-terminal sequence. Sequences between zinc fingers that were constrained among vertebrates did not share strong sequence similarity across invertebrate lineages, except for the C4-like region. While broadly distributed, ZNF423 homologs appeared to be absent from some well-annotated genomes, including Ciona and other urochordates, several Caenorhabditis species, and animals outside of Bilateria.

Invertebrate homologs had lineage-specific C2H2 ZF number and identity, although available sequence accessions were dominated by insects (Figure 2 and Supplemental Table S2). While gene models should be considered provisional, models that began with the conserved MSRRKQ and had well-defined carboxy-terminal sequences (e.g., strong matches to vertebrate ZF28-30 or coherence within a larger invertebrate clade) are probably approximately correct. Comparatively deep sampling of Arthropod sequences supported the idea that structural differences in predicted protein sequences primarily reflected lineage-specific changes rather than annotation errors and, at least within insects, changes in number of C2H2 fingers occurred primarily between orders rather than other taxonomic units. Excluding gene models that appeared to be truncated annotations (missing terminal exons relative to other members of their taxonomic groups), typical arrangements showed 21 ZFs in Diptera (11 of 12 gene models from distinct genera and each of several Drosophila species), 22 ZFs in Coleoptera (4 of 6), and 24 ZFs in Hymenoptera (18 of 23) and Hemiptera (7 of 10) and 28-29 ZFs in Blattodea (3 species). Hymenoptera also showed a larger ZF12 (44 amino acids, position corresponding to human ZF18) than other insect groups. Further diversity was seen in the few sequences available from other Arthropod orders (24-36 ZF among arachnids, 35 in horseshoe crab).

**Figure 2.**
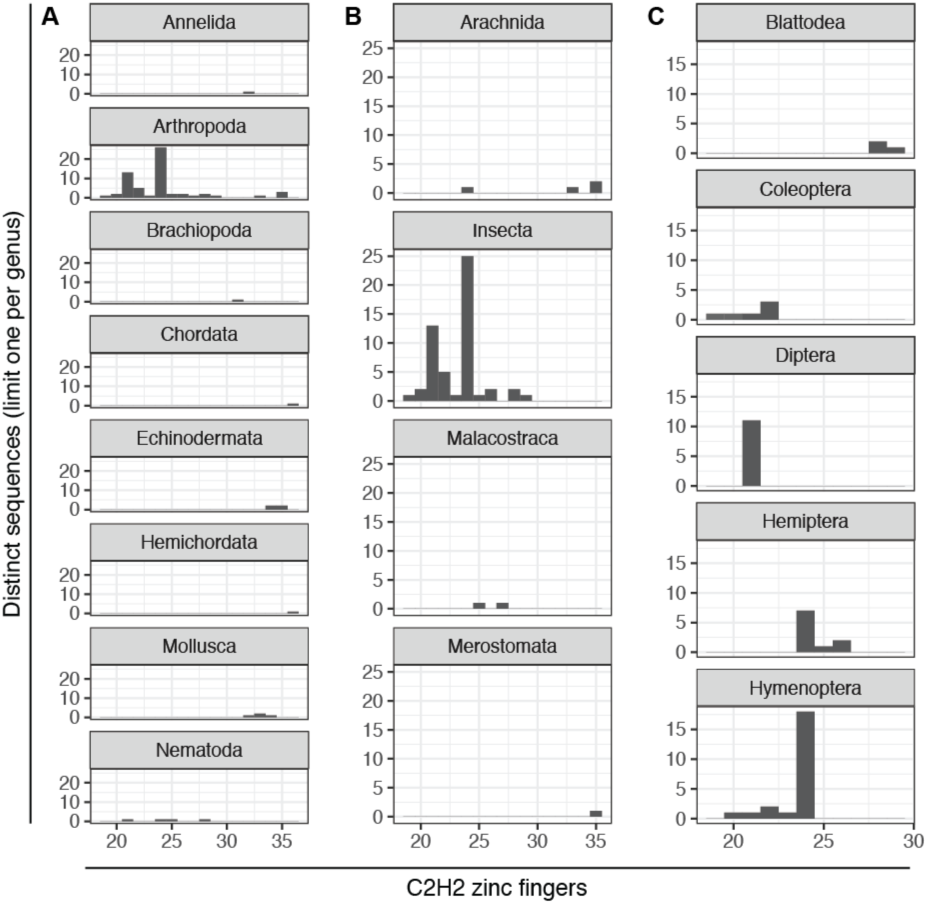
Distribution of C2H2 ZF number among invertebrate ZNF423 homologs shows remodeling over longer timescales. (A) Invertebrate phyla show differences in distribution. Far greater number of sequences available among arthropods allows comparisons at finer scales. (B) Among arthropod classes, arachnids had more ZFs, while most of the available sequences were from insects. (C) Insects orders had characteristic modal ZF number. Only one accession per genus was included.

### Differences in zinc finger number include both terminal and internal changes

Because of the repeated-C2H2 ZF structure of ZNF423 homologs and repeated changes in ZF number, long-range alignments might shift between homologous and non-homologous fingers. To more clearly define individual ZF homologies, I used two approaches: pair-wise reciprocal best BLAST matches between individual fingers of two homologs and multiple sequence alignment trees for isolated ZFs from diverse ZNF423 homologs, and compared to their position of origin for subsets of ZNF423 homologs. Although both approaches suffer from the limited information content and differentiation among 22-26 aa C2H2 domains, in which several residues are either invariant (4 zinc-coordinating C and H positions) or highly constrained (3 hydrophobic positions that form the core of the ZF fold) among ZFs, both approaches showed strong conservation of ZF order and cluster identity among the most-conserved ZFs.

Several homologies were consistent across large taxonomic distances. ZFs homologous to the human GCACCC DNA or SMAD protein binding clusters were typically the most constrained sequences, followed by EBF-binding ZFs (Figures 3 and 4). Changes in ZF number were evident both at the ends of the protein and within homologous clusters, seen more readily from mapped pairwise comparisons in Figure 3. Invertebrate deuterostomes typically included a range of additional C-terminal ZFs beyond fingers homologous to the EBF-binding cluster in humans (Figure 3A). These extended domains tended to cluster in multiple sequence alignments, but were generally less conserved than functionally annotated ZFs and many of the novel C-terminal ZFs clustered as outer branches with weak similarity. The C4-like sequence noted from vertebrate alignments occupied the same position relative to the EBF-associated C2H2 cluster across species, including those where the homologous region met criteria for a RING domain.

**Figure 3.**
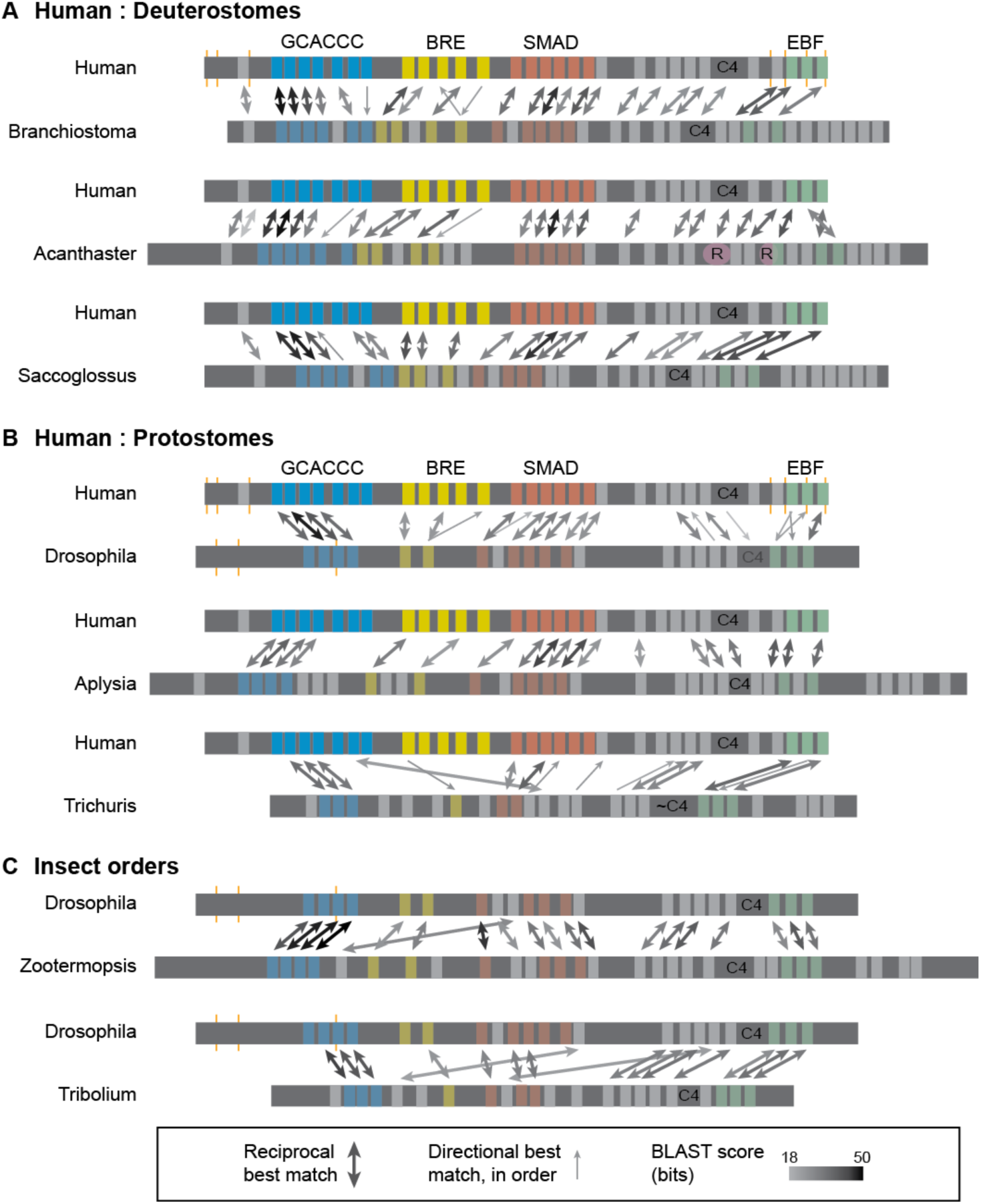
Inferred homologies of distinct C2H2 fingers between human and invertebrate homologs. BLAST alignment maps from full-length protein query and libraries of each C2H2 domain of a target homolog (light grey and colored stripes). Double arrows are reciprocal best matches, with shading scaled to the average score for reciprocal comparisons. Thin single arrows are best matches in only the indicated direction, but conform to approximate domain order. ZFs without arrows did not have a match that met either criterion. (A) Comparisons between human ZNF423 and invertebrate deuterostomes, included a chordate (Branchiostoma), an echinoderm (Acanthaster) and a hemichordate (Saccoglossus). Human ZNF423 C2H2 clusters associated with binding to GCACCC DNA sequences, BMP response element (BRE), SMAD proteins and EBF-family proteins are indicated, with corresponding fingers color coded. Inferred cognates in target species are similarly colored, with reduced intensity. Position C4 ZF-like sequence, which in Acanthaster corresponded to a RING (R) domain, are indicated in each homolog. (B) Comparisons between human ZNF423 and three protostomes, including an arthropod (Drosophila), a mollusk (Aplysia) and a nematode (Trichuris). (C) Changes in ZF number between insect orders.

**Figure 4.**
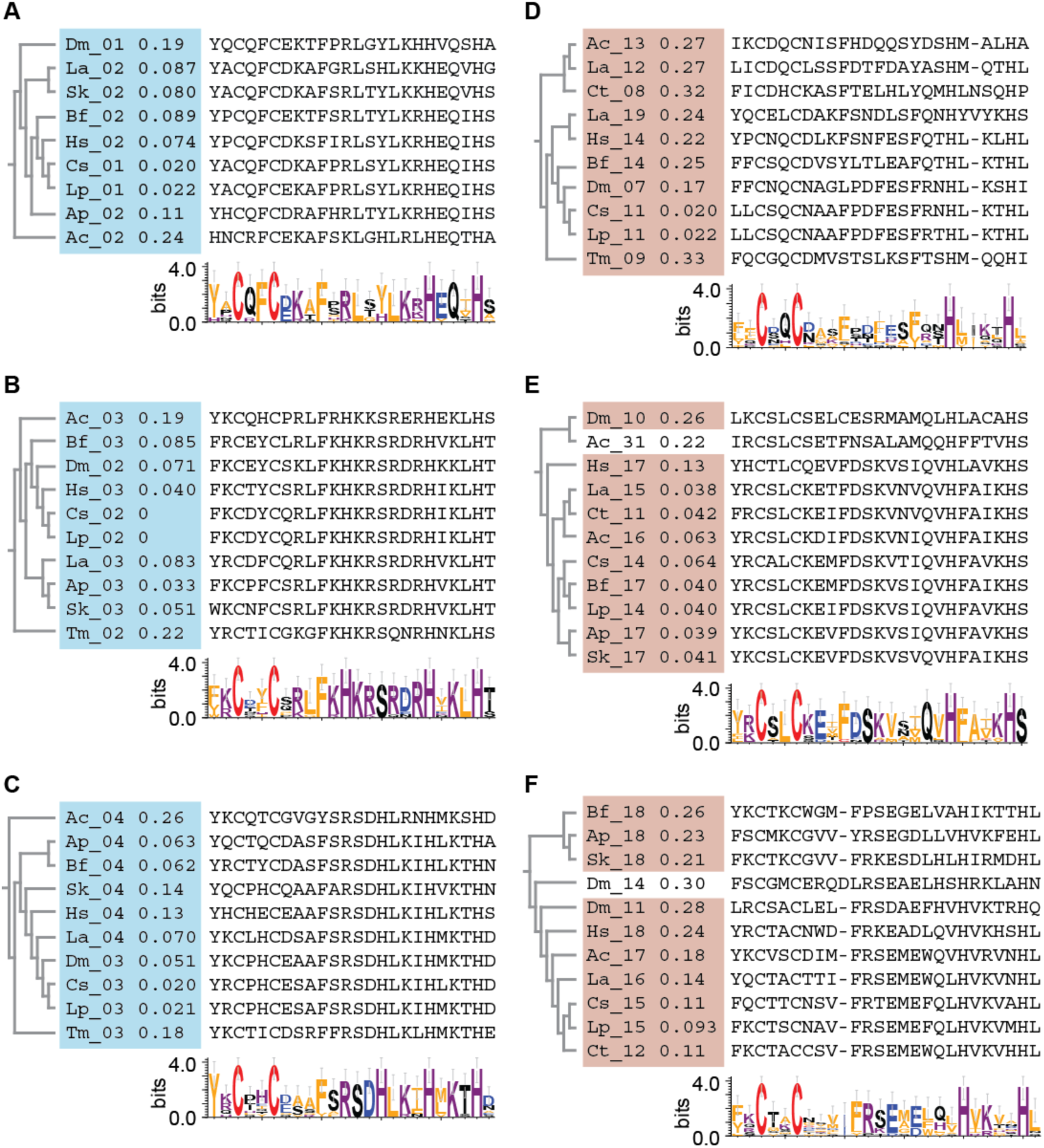
Multiple sequence alignment clusters C2H2 ZFs by position. Sub-trees within the alignment of all ZFs in 11 ZNF423 homologs are shown. Sub-trees were selected around highly conserved and functionally annotated human ZF sequences from the GCACCC DNA-binding (blue shading, A-C) or SMAD-binding (light brown shading, D-F). Sequence alignments were simplified within each sub-tree to remove shared gaps created in the larger alignment. Differences within sub-trees were few and detailed topologies should be interpreted cautiously. Even for well-conserved fingers, not all species aligned a ZF within a given sub-tree and additional ZFs (Ac_31, Dm_14) invade the tree. Sequence logo height is scaled to information content of residues at a given position. Sequences were from human (Hs, Chordata, subphylum Craniatata), *Branchiostoma floridae* (Bf, Chordata, subphylum Cephalochordata), *Acanthaster planci* (Ap, Echinodermata), *Saccoglossus kowalevskii* (Sk, Hemichordata), *Aplysia californica* (Ac, Mollusca), *Lingula anatina* (La, Brachiopoda), *Capitella teleta* (Ct, Annelida), *Drosophila melanogaster* (Dm, Arthropoda, class Insecta), *Centruroides sculpturatus* (Cs, Arthropoda, class Arachnida) *Limulus polyphemus* (Lp, Arthropoda, class Merostomata), and *Trichuris muris* (Tm, Nematoda) predicted by SMART with outliers allowed and manually reviewed.

Changes in ZF number were more pronounced in protostome homologs, including changes within annotated clusters encoded by a single large exon in both vertebrate and Diptera genomes (Figure 3B). For example, the Drosophila homolog appeared to have fewer ZFs in both the GCACCC-binding and BRE-binding homologous clusters, as did a Trichuris nematode homolog. This was also true among insect orders with characteristic, but different, number of C2H2 ZFs (Figure 3C). The termite *Zootermopsis nevadensi*s (29 ZF, order Blattodea) had both more ZFs among internal clusters and four additional ZFs C-terminal to the EBF-related cluster relative to *Drosophila melanogaster* (21 ZF, order Diptera). The beetle *Tribolium castaneum* (24 ZF, order Coleoptera) had more ZFs internally relative to Drosophila, but also terminated with the EBF-related cluster.

To complement pair-wise analysis, each ZF from 11 species (four deuterostomes and seven protostomes) representing 8 phyla were used for multiple sequence alignment and assessed for both overall tree structure and amino acid-level conservation. Neighbor-joining trees from either MUSCLE or MAFFT alignments produced sub-trees of ZFs from similar positions in their respective proteins, generally confirming position-specific homology. Some species left outside the position-specific branches and some ZFs from different positions invading the branches, even for the highly conserved ZFs in the DNA-binding (Figure 4A-C) and SMAD-binding (Figure 4D-F) clusters, possibly due to the limited information content in short peptides. For even the most-conserved fingers, no sub-tree included a unique ZF for all 11 species before encountering a second ZF from another position in at least one species. Thus, while the limited information content in any one ZF sequence supports homology of position-specific ZFs generally, homology among ZFs with less robust sequence identity may be better inferred by accounting for context of position within the ZF clusters and contiguous homology of adjacent ZFs.

### Zinc fingers beyond 30 were ancestral and lost in vertebrates

Both deuterostome and protostome lineages have examples with C2H2 ZFs C-terminal to those that align with vertebrate ZFs. ZF-specific alignments support homology between deuterostome and protostome C-terminal ZFs, suggesting that they represent an ancestral state independently lost in vertebrate and insect lineages. Pair-wise alignment between the chordate *Branchiostoma floridae* and the mollusk *Aplysia californica* illustrates that best reciprocal matches between isolated ZF sequences maintains coherent order through the C-terminal domains (Figure 5A). Sub-trees from an alignment of all fingers from the same 11 species used in Figure 4 supported homology among invertebrates for ZFs aligned to Branchiostoma ZF30 (Figure 5B), ZF31 (Figure 5C) ZF32 (Figure 5D) and ZF33-ZF35 (Figure 5E). ZF domains that did not fall within the sub-trees predicted from position show similarity beyond those required by the ZF domain definition and may highlight specific residues or properties selected by evolution that are missed by simple alignment (Figure 5B-D). Among the 11 species in the all-ZF alignment, nine had C-terminal extensions relative to human and Drosophila. Each of these nine had at least two ZFs that fell within sub-trees that corresponded by position (Figure 5F). This supports the idea of additional binding characteristics for this homology group that are conserved among many species outside of vertebrates and insects.

**Figure 5.**
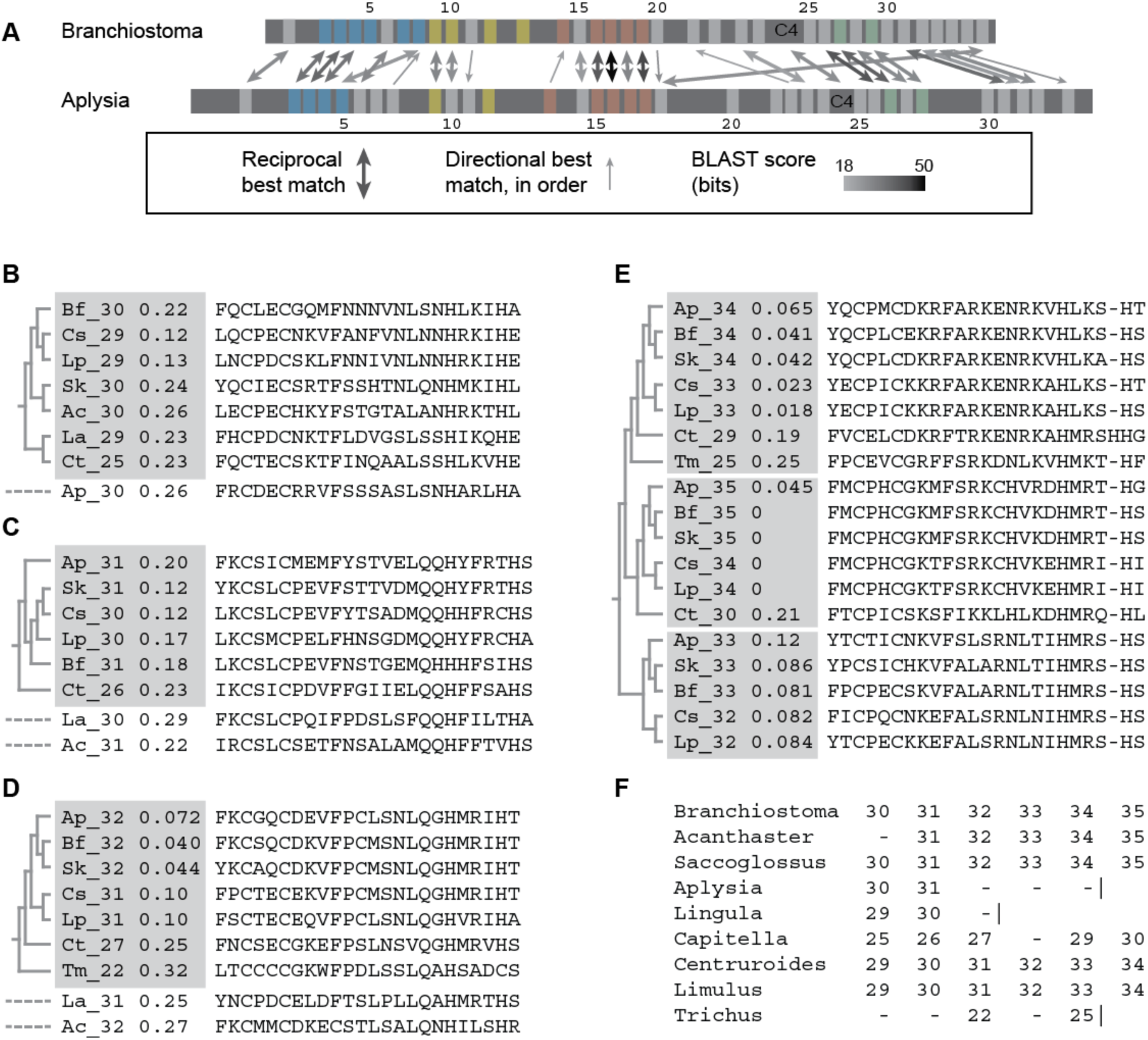
Zinc fingers C-terminal to vertebrate alignments share homology across deuterostome and protostome lineages. (A) Best reciprocal matches between ZF domains of deuterostome *Branchiostoma floridae* and protostome *Aplysia californica*. Double arrows are reciprocal best matches, with shading scaled to the average score for reciprocal comparisons. Thin single arrows are best matches in only the indicated direction, but conform to approximate domain order. ZFs without arrows did not have a match that met either criterion. (B-E) Sub-trees from MAFFT alignment of all ZFs from the 11 diverse species listed in Figure 4 cluster by relative position and support homology for 6 C-terminal ZFs. (B-D) For the first three positions, ZFs from the same relative position that did not fall within the sub-tree are indicated with a dashed line below the tree. (E) Clusters for the next three positions are adjacent in the all-ZF tree. (F) Each of the nine species with C-terminal ZFs included at least two that aligned by position. Dashed indicate non-aligned ZFs, vertical lines indicate end of the protein after that ZF. Human and Drosophila were included in the alignment but did not contribute ZFs to these sub-trees. Bf, *Branchiostoma floridae*; Ap, *Acanthaster planci*; Sk, *Saccoglossus kowalevskii*; Ac, *Aplysia californica*; La, *Lingula anatina*; Ct, *Capitella teleta*; Cs, *Centruroides sculpturatus*; Lp, *Limulus polyphemus*; *Tm, Trichuris muris*.

### Domain-level homology includes the C4 zinc finger-like segment

Reciprocal alignments of annotated human ZNF423 ZF domains to invertebrate homologs (and invertebrate fingers to human ZNF423) showed conserved order of ZFs and ZF clusters as a general feature, changes in number notwithstanding (Figure 3). This analysis further supported conservation of the C4-like sequence, which retained a constant position relative to SMAD-related and EBF-related clusters (equivalent to sequence between human ZF25-ZF26), rather than to overall C2H2 number, across all lineages examined (Figure 3). In a few homologs, such as the sea star Acanthaster, the C4-like sequence was expanded and met annotation criteria for a RING domain, while remaining a reciprocal best match to the human C4-like region.

Within the C4 -like sequence, the CxxC sites were separated by a range of 10 (several aphid species) to 31 (termites and cockroach) residues in all homologs. As vertebrate homologs showed little sequence variation throughout the full-length protein (Figure 1A), analysis of constraints on this C4 zinc finger region focused on invertebrate homologs (nine Drosophila species, mutually separated by ≥20 M years were added for this analysis, to accommodate order-level divisions below). The five invertebrate homologs that did not have two CxxC motifs included two arthropods with a looser configuration of 5 Cys (Hyallella) or a CxxC plus several His (Armadillidium) residues, two nematodes with a single CxxC plus other Cys residues (Trichinella and Trichuris) and one nematode with a RING domain (Brugia).

Deep coverage of Arthropod species again illustrated the dynamics for this putative domain. MAPP analysis showed constraint in the C4-like sequence approaching that of C2H2 domains within both Diptera (Figure 6A) and Hymenoptera (Figure 6B). However, unlike C2H2 ZFs the characteristic cysteine residues in Diptera (Figure 6C) and Hymenoptera (Figure 6D) stand out in constraint relative to neighboring positions. Extending this to 58 Arthropod sequences showed very limited information content from other positions in a MAFFT alignment (Figure 6E).

**Figure 6.**
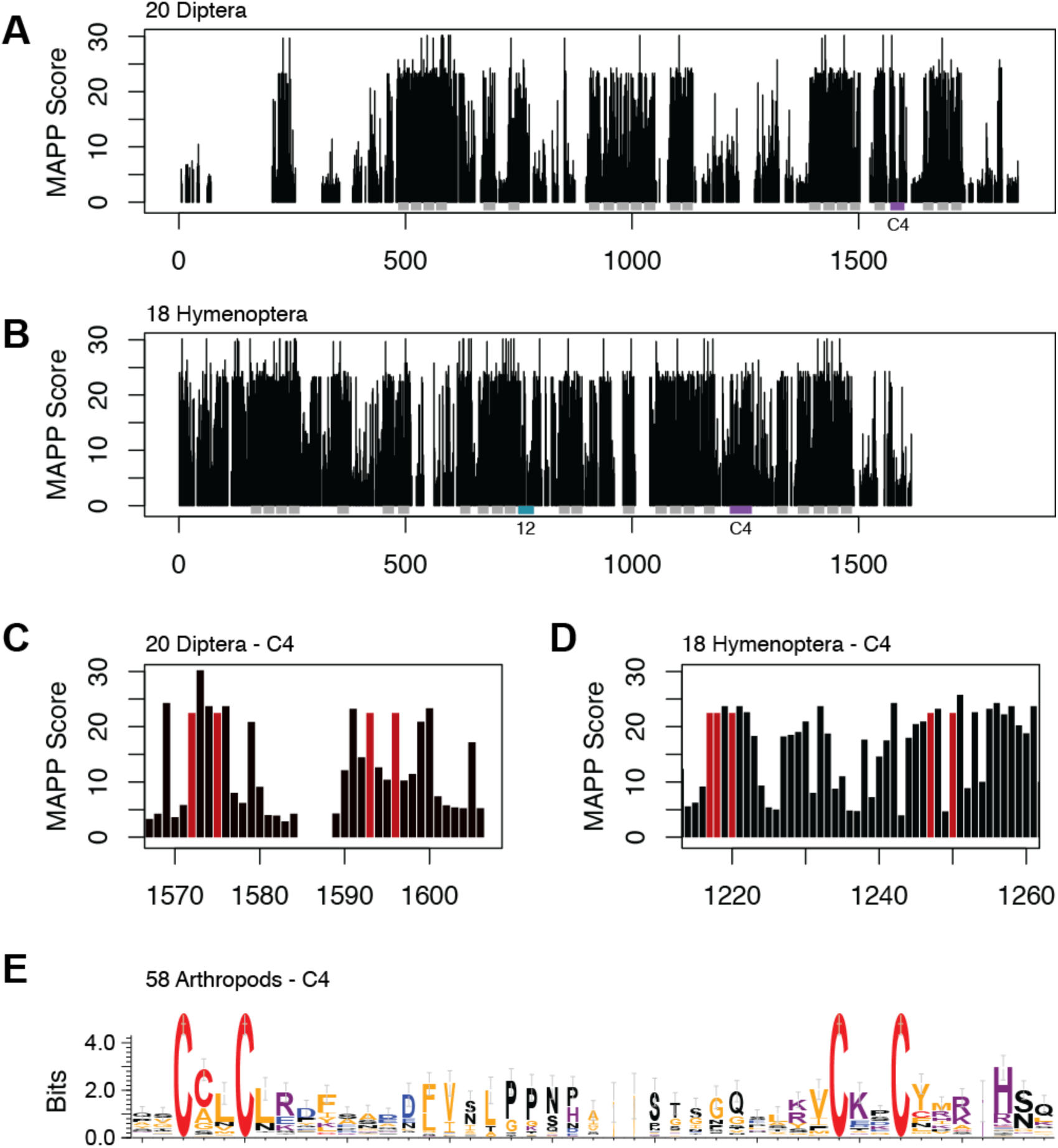
The C4-like region is a conserved feature with limited sequence constraint. (A) MAPP analysis of 20 aligned Dipteran homologs from different genera show strong constraint within C2H2 domains (grey bars). The putative C4 region (purple) shows a gap in MAPP score due to variability in the length between the two CxxC motifs. (B) MAPP analysis for 18 Hymenopteran homologs, where CCxC to CxxC spacing is less heterogeneous shows constraint near that of C2H2 domains. The enlarged ZF12 is highlighted. (C,D) MAPP analysis showing just the C4-like sequences of Diptera (C) and Hymenoptera (D) with absolutely conserved cysteine residues in red. (E) WebLogo representation of MAFFT-alignment for C4-like region of shows complete conservation of CxxC motifs and much less constraint at other residues across 58 Arthropod ZNF423 homologs.

## Discussion

The multiple zinc finger clusters in ZNF423 provide a physical scaffold for multiple protein partners and site specific DNA binding. Prior observations that ZNF423 is prominently expressed in immature precursor cells and that its partner proteins have mutually inhibitory relationships supports a view that ZNF423 serves in part to integrate signaling pathways during developmental programs. Depletion of loss-of-function variants in human genetic databases as well as structural abnormalities in animal models support the idea that ZNF423 is critical for developmental processes in brain and other tissues. This paper examined conserved features among inferred ZNF423 orthologs and levels of constraint across several animal taxa and identified several features not previously identified from vertebrate or Drosophila homologs. This extended analysis identified a candidate C4-class ZF not previously identified in this well-annotated protein family and demonstrated additional C2H2 fingers that are conserved among several invertebrate lineages.

While vertebrate homologs showed very strong sequence constraint across the entire protein, an expanded set of invertebrate homologs showed extensive remodeling of ZF number. Changes in ZF number occurred both within clusters that were homologous to vertebrate ZF clusters known to binding DNA, SMADs, and EBFs. Direct inference of domain-level orthology among specific ZFs is inherently limited by the small size (22-27 aa) of C2H2 ZFs, but inferences drawn here were supported by consistent conservation of order and clustering across several pair-wise comparisons. Zinc fingers that extended beyond homology to vertebrate ZNF423 homologs were nonetheless homologous to each other, even between deuterostome and protostome lineages, separated ∼800 Mya, suggesting that lack of these fingers in both vertebrates and related orders of insects represent independent loss events. Rather than the very static view from vertebrate ZNF423 constraints, the expanded set of homologs suggest a view of ZNF423 as a modular and adaptive platform for integrating developmental signals across transitions in developmental plans of many animal lineages. Whether apparently homologous ZF clusters bind to orthologous targets and what new targets might be specified by reconfigured or novel ZF clusters them remains to be determined. Constraint scores shown in this paper should be interpreted with caution as selection of species for analysis was inherently biased by the availability and quality of sequenced genomes. Deeper sampling of currently sparse or absent lineages will likely refine the estimated distribution of ZF number and placement of variant C2H2 fingers, as well as constraints on specific residues.

The analysis here identified a potential C4-like ZF that has not been included in previous annotations owing to the limited sequence constraint for this class of zinc finger. By comparing C2H2 ZF similarities, the analysis here showed that a C4-like sequence (a full RING finger in three species) is a conserved feature of ZNF423 homologs. The same feature is seen among vertebrate ZNF521 paralogs. While Tsai and Reed’s original observation that this sequence may contribute to ZNF423 multimer formation [15], whether this sequence feature has a consistent structure, whether it binds zinc, and what function it imparts to the protein remain to be discovered.

## Methods

### Identification of ZNF423 homologs

Protein sequences were identified by iterative and reciprocal BLASTP and TBLASTN [44-46] searches of public databases and from curated orthologs in Metazome (https://metazome.jgi.doe.gov/) and OrthoDB [47], https://www.orthodb.org/). BLAST searches were conducted using the NCBI (https://blast.ncbi.nlm.nih.gov/Blast.cgi) and the EBI BLAST (https://www.ebi.ac.uk/Tools/sss/ncbiblast/) web interfaces. *Lytechinus variegatus* and *Patiria miniata* homologs were obtained from EchinoBase ([48] http://www.echinobase.org/Echinobase/). Taxonomy-delimited searches were done as a final step to identify potential homologs in sparsely-covered or absent lineages. For each genus, the best reciprocal match was considered first. After identification of the conserved MSRRK N-terminal sequence as taxonomically more wide-spread than the N-terminal sequence of the human RefSeq protein, gene models that contained MSRRK where multiple gene models were found within a genus. Sequences denoted low-quality in their annotation or containing ambiguity positions were excluded. Sequences shorter than 1000 aa were not considered for most analyses to avoid truncated annotations.

### Sequence analyses

Domain annotations used SMART ([49, 50], http://smart.embl.de/) with manual review and cross-validation of selected examples in InterPro [51, 52], https://www.ebi.ac.uk/interpro/). For alignment and manual curation C2H2 ZFs were considered to include two residues before the first cysteine and one residue after the second histidine. Alignments were performed with default parameters in MUSCLE [53, 54] https://www.ebi.ac.uk/Tools/msa/muscle/) or MAFFT [55] using the EMBL-EBI web interface [56]. Physico-chemical constraints were assessed in MAPP [40], downloaded from http://mendel.stanford.edu/sidowlab/downloads/MAPP/index.html and run in Java under MacOS 10.14.3. Median scores among all possible substitutions at each position (column score) were plotted as a histogram in R 3.5.1. For non-synonymous to synonymous substitution ratio (dN/dS), corresponding cDNA sequences were aligned at the codon level using PAL2NAL [57] (http://www.bork.embl.de/pal2nal/) and dN/dS calculated by single likelihood ancestor counting using the SLAC tool in Datamonkey [58, 59] from its web interface (http://www.datamonkey.org/). For comparison and cross-validation with different measures of constraint, similar analysis on vertebrate alignments were performed from web interfaces for ConSurf (https://consurf.tau.ac.il/) and ScoreCons (https://www.ebi.ac.uk/thornton-srv/databases/cgi-bin/valdar/scorecons_server.pl) using default parameters [41, 42]. Sequence logo displays were created in WebLogo 3.7.4 ([60] http://weblogo.threeplusone.com/) using the indicated alignments and a custom color scheme to highlight the conserved cysteine residues in predicted C2H2 and C4 zinc fingers.

## Acknowledgements

I thank Amit Majithia, Kevin Ross, and Arend Sidow for helpful comments on a draft manuscript. This work was supported by grant R01 NS097534 from the National Institute of Neurological Disorders and Stroke.

